# The application of community ecology theory to co-infections in wildlife hosts

**DOI:** 10.1101/2020.04.15.042937

**Authors:** Chloe Ramsay, Jason R. Rohr

## Abstract

Priority effect theory, a foundational concept from community ecology, states that the order and timing of species arrival during species assembly can affect species composition. Although this theory has been applied to co-infecting parasite species, it has almost always been with a single time lag between co-infecting parasites. Thus, how the timing of parasite species arrival affects co-infections and disease remains poorly understood. To address this gap in the literature, we exposed post-metamorphic Cuban tree frogs (*Osteopilus septentrionalis*) to Ranavirus, the fungus *Batrachochytrium dendrobatidis* (Bd), a nematode *Aplectana hamatospicula*, or pairs of these parasites either simultaneously or sequentially at a range of time lags and quantified load of the secondary parasite and host growth, survival and parasite tolerance. Prior exposure to Bd or *A. hamatospicula* significantly increased viral loads relative to hosts singly infected with Ranavirus, whereas *A. hamatospicula* loads in hosts were higher when co-exposed to Bd than when co-exposed to Ranavirus. There was a significant positive relationship between time since Ranavirus infection and Bd load, and prior exposure to *A. hamatospicula* decreased Bd loads compared to simultaneous co-infection with these parasites. Infections with Bd and Ranavirus either singly or in co-infections decreased host growth and survival. This research reveals that time lags between co-infections can affect parasite loads, in line with priority effects theory. As co-infections in the field are unlikely to be simultaneous, an understanding of when co-infections are impacted by time lags between parasite exposures may play a major role in controlling problematic co-infections.

## Introduction

Simultaneous infection with multiple species of parasites, hereafter referred to as co-infection, is a regular occurrence for host species (Greub et al. 2000, Kim et al. 2003, Swanson 2006). Direct interactions among these parasites can occur when each occupies similar habitats in hosts, competing for space or resources and, in some cases, they can even depredate one another (Kuris and Lafferty 1994, de Roode et al. 2004). Indirect interactions of parasites can also occur. These can be facilitative or inhibitory and are often mediated by the host immune system. As an example, microparasites generally activate the T helper 1 (Th1) arm of the adaptive immune system (fungal infections can also activate Th17; Berger 2000, Blanco and Garcia 2008, Zelante et al. 2009), whereas macroparasites activate the Th2 arm of the immune system (Paul and Zhu 2010). Because an investment in one arm generally comes at the expense of investment in the other (i.e. a tradeoff; Abbas et al. 1996, Morel and Oriss 1998, Yazdanbakhsh et al. 2002), micro-marcroparasite co-infections are often facilitative (Abbas et al. 1996, Berger 2000), whereas micro-microparasite and macro-macroparasite co-infections can be inhibitory though cross-immunity (Johnson and Buller 2011).

Interactions among parasites within hosts can affect the ability of hosts to tolerate infections (e.g. a host’s ability to maintain high health despite high infection loads; Rohr et al. 2010) and thus can have important consequences on disease dynamics, host fitness, and population growth rates (Corbett et al. 2003, Bromenshenk et al. 2010, Ezenwa and Jolles 2015). For example, co-infections with multiple virus strains and microsporidia were detected in honeybee hives suffering from colony collapse disorder (Bromenshenk et al. 2010). Deworming a population of African buffalo increased the transmission of the deadlier *Mycobacterium bovis* (Ezenwa and Jolles 2015). Mice co-infected with influenza virus and *Legionella pneumophila* were less tolerant of infection than singly infected mice, increasing their mortality (Jamieson et al. 2013).

Understanding how co-infecting parasites interact with one another and their host could be informed by community ecology theory which highlights interactions among species and their environment (Pedersen and Fenton 2007, Johnson et al. 2015). Priority effects in ecology describe how the order and timing of species arrival during species assembly affect species composition (Sutherland 1974, Drake 1991, Leopold et al. 2017). In community ecology, priority effects have been shown to affect predator-prey interactions (Blaustein and Margalit 1996), competition (Wilbur and Alford 1985, Sousa 1992), and community succession (Facelli and Facelli 1993, Devevey et al. 2015, Leopold et al. 2017). Priority effects have also been shown to be important in parasite-parasite interactions (Sousa 1992, de Roode et al. 2005, Hoverman et al. 2013, Devevey et al. 2015). For example, relative to simultaneous co-infections, infection with one strain of *Plasmodium chabaudi*, a causative agent of malaria, reduced infections from a second strain of the pathogen when challenged three days after the first (de Roode et al. 2005).

The definition of priority effects encompasses both order and timing of species arrival (Sutherland 1974, Drake 1991, Leopold et al. 2017). However, most studies applying priority effect theory in co-infecting parasite species have focused exclusively on the order and thus have had a single time lag between co-infecting parasites (Hoverman et al. 2013, Devevey et al. 2015, Leopold et al. 2017). Thus, how the timing of parasite species arrival affecting co-infections and disease remains poorly understood (but see de Roode et al. 2005, Wuerthner et al. 2017 for three time points). This is an important gap in the literature because co-infections are almost always staggered in nature, making it important to understand how time lags between co-infections affect disease. In community ecology, the initial population size of interacting species can affect the outcome of their competition, with larger initial populations being more likely to outcompete and potentially competitively exclude an introduced second species (Chesson and Kuang 2008). Therefore, for microparasites that internally replicate and are not easily cleared, longer time lags should allow the first parasite to reach higher densities in the host before a second parasite attempts to infect (Rojas et al. 2005, Rosenblum et al. 2012). Additionally, longer time lags and higher microparasite densities would allow for and cause elevated immune responses, respectively, increasing the likelihood that immunity alters co-infection dynamics (Graham 2008, Piecyk et al. 2019).

The outcomes of co-infections are particularly important for amphibians because amphibians are suffering from worldwide declines and extinctions due in large part to the spread of several diseases. As a result, amphibians have become the most threatened vertebrate taxon on the planet, with 32.5% of species threatened globally (Stuart et al. 2004, Wake and Vredenburg 2008, Scheele et al. 2019). The primary goals of this study are to test how a co-infecting parasite and the time lag between co-infections affects the secondary parasite and amphibian host fitness (growth and survival) and whether these interactions are mediated by host immune responses (indirect effects). To address these goals, we exposed Cuban treefrogs (*Osteopilus septentrionalis*) to all pairwise combinations of a fungus (microparasite), virus (microparasite), and nematode (macroparasite). The fungus we studied is *Batrachochytrium dendrobatidis* (Bd), a chytrid fungus associated with hundreds of amphibian declines and extinctions globally (Berger et al. 1998, Skerratt et al. 2007, Kilpatrick et al. 2010). This fungus causes the disease chytridiomycosis, which leads to cardiac arrest by fusing keratinous portions of amphibian skin and thus, limiting electrolyte transport (Voyles et al. 2009, Kilpatrick et al. 2010). The virus we studied is frog virus 3 (FV3), a Ranavirus from the family *Iridoviridae* that replicates in the internal organs of ectothermic vertebrates (Chinchar 2002), causing hemorrhaging and mass mortality (Gantress et al. 2003, Gray et al. 2009). FV3 is also associated with mass die-offs of amphibians across the planet (Green et al. 2002, Gray et al. 2009). In addition to these deadly microparasites, treefrogs were also exposed to a macroparasite, specifically the gastrointestinal nematode, *Aplectana hamatospicula*. This nematode is a common parasite of amphibians in the Southeastern US and Latin America (Baker 1987, Vhora and Bolek 2013, Ortega et al. 2015). Importantly, parasitic worms, Bd, and Ranaviruses are known to co-occur in amphibian hosts in the field (Hoverman et al. 2012, Stutz et al. 2018, Watters et al. 2018). In the Peruvian Andes, an amphibian biodiversity hotspot, up to 50% of frogs were coinfected with Bd and Ranavirus (Warne et al. 2016).

Our goals were motivated by several hypotheses. First, the three focal parasites occupy different microhabitats within the host; Bd infects the skin, *A. hamatospicula* dwells in the gastrointestinal tract, and Ranavirus primarily infects the kidneys and liver (Cunningham et al. 1996, Voyles et al. 2009, Knutie et al. 2017). Thus, we hypothesized that these parasites would primarily interact indirectly, mediated by host immunity. Second, because microparasites activate the Th1 arm and macroparasites activate the Th2 arm of the host immune system (Abbas et al. 1996) and investment in each arm is mutually exclusive (Graham 2008), we hypothesized that co-infections that involve any microparasite (Bd and Ranavirus) and the macroparasite (*A. hamatospicula*) would be facilitative. In contrast, we hypothesized that Bd and Ranavirus co-infections might be inhibitory because they activate the same arm of the immune system. Third, we hypothesized that longer time lags between initial and secondary infection would increase the strength of co-infection priority effects for micro-but not macroparasites. This is because microparasites, but not macroparasites, can replicate in the host and thus greater time lags would allow for greater time for microparasite densities to increase. In this study, a priority effect is defined as any significant relationship between the time lag between exposure to the first and second parasite and the load of the second parasite. Fourth, from a host health perspective, we hypothesized that host growth and survival would be lower if the initial parasite *i*) was particularly virulent relative to the other co-infecting parasite and *ii*) had a long time to replicate within the host, reaching high parasite loads (assuming it was a microparasite). Fifth, when co-infecting parasites inhibit one another (either directly or through the host immune system), we expected co-infection to increase host tolerance (e.g. a host’s ability to maintain high health despite high infection loads), whereas when co-infecting parasites were facilitative of one another, we hypothesized co-infection to decrease host tolerance.

## Methods

### Animal Husbandry

Post-metamorphic Cuban treefrogs (*Osteopilus septentrionalis*) were collected from PVC pipes (7.62 x 91.44 cm) hammered into the ground upright around wetlands in Flatwoods Wilderness Park, Tampa, Florida (28.11° N, 82.31° W) from February to March 2016. Individuals were housed separately in 57.75 cm^3^ sterilized plastic cups containing an unbleached paper towel wet with artificial spring water (ASW). Tree frogs were fed 2-week-old crickets *ad libitum* twice a week at 25 °C (12hr photoperiod). Each of these plastic cups was washed by hand, bleached in a 10% solution, thoroughly rinsed with deionized (DI) water, and dried weekly.

### Pre-inoculation

To eliminate any potential helminth infections in wild-caught frogs, collected individuals were orally treated with ivermectin (0.2 mg/kg of host) two months before the experiment. Following ivermectin exposures, husbandry containers were changed daily for one week to reduce the potential for reinfection with any larval *A. hamatospicula*. Once frogs did not shed helminths for four weeks, they were considered uninfected. None of our control animals tested positive for Bd or Ranavirus throughout the course of the experiment, suggesting that these amphibians were not infected when they were collected, nor did they become inadvertently infected during the experiment. Wild-caught frogs were randomized into treatment groups so that treatment groups all had similar average frog weights, evenly distributing the larger and smaller frogs. Two days before initial parasite exposure, frogs were moved to an environmental chamber to acclimate to experimental conditions. The environmental chamber, where amphibians remained for the duration of the experiment, was kept at 17 °C (12h photoperiod) because Cuban treefrogs can clear Bd at higher temperatures and 17 °C is an intermediate temperature for Ranaviral growth *in vitro* (Chinchar 2002, McMahon et al. 2014, Cohen et al. 2017, Price et al. 2019).

### Parasite culture

*A. hamatospicula* were obtained from the gastrointestinal tract of euthanized Cuban treefrogs collected from Flatwoods Wilderness Park. Extracted worms were stored in a Petri dish (15 x 150 mm; *n* ≤ 5 per dish) filled with 2mL of DI water and held under dark conditions. We collected larval worms (J1) from adults and held each for ≥ 14 days to ensure maturation to the infectious stage (*i.e*., J3). Gastrointestinal (GI) content was collected from uninfected frogs and stored identically to act as a sham inoculant.

Ranavirus was isolated from infected wood frogs (*Lithobates sylvaticus*) in Michigan and has a 99% similarity to a ∼500 bp fragment of the MCP gene of *Frog Virus 3* (GenBank Accession number: PRJNA504607). The virus was cultured in fathead minnow cells (*Pimephales promelas*) and stored in minimal essential media (MEM) at -80 °C.

Bd (strain SRS-JEL 212) solution was grown on 1% tryptone-agar Petri dishes, aliquoted from an active culture. Bd growth occurred over eight days at 23°C until zoospores were observed. Petri dishes were then flooded with 1 mL of DI water for five minutes to permit zoosporangia to release infectious zoospores. Released zoospores and DI water were then poured into a beaker with 15 mL of DI water and stirred to produce a homogeneous slurry. To assess Bd zoospore concentration, we sampled the homogeneous slurry and counted zoospore density using a hemocytometer. We added DI water to the slurry to obtain a final Bd concentration of 10^5^ zoospores/ml. Sterile 1% tryptone plates without Bd were cultured at 23°C for eight days and rinsed with DI water to produce sham Bd and control treatments.

### Experimental Design

To examine how the timing of initial parasite exposures affects host parasite dynamics, we exposed Cuban treefrogs to one of ten parasite treatments. Parasite treatments included exposure to *A. hamatospicula* alone, Ranavirus alone, or Bd alone, exposure to pairwise combinations of two parasites in sequential order (six co-infection treatments), and a sham exposure (control; see below for details). Each of the six co-infection treatments had five temporal subtreatments of staggered exposures (Table 1). Five frogs were assigned to each of the 34 treatment combinations for a total of 170 experimental individuals.

**Table 1.** An example of the general design of the timing of the initial and secondary infection are shown below for one pairwise combination of parasites, single infections, and controls. The same general design was applied to all other pairwise combinations of parasites. There were 5 individuals for each treatment group. Bd = *Bactrachochytrium dendrobatidis*, Rv = Ranavirus.

|  | Proportion of time lags (time lag : the maximum time lag) |  |  |  |  |
| --- | --- | --- | --- | --- | --- |
| Exposures | -1 to -0.76 | -0.75 to -.51 | -0.5 to -0.26 | -0.25 to -0.01 | 0 |
| Bd |  |  |  |  | Bd |
| Rv then Bd | Rv |  |  |  | Bd |
|  |  | Rv |  |  | Bd |
|  |  |  | Rv |  | Bd |
|  |  |  |  | Rv | Bd |
|  |  |  |  |  | Rv & Bd |
| Rv |  |  |  |  | Rv |
| Bd then Rv | Bd |  |  |  | Rv |
|  |  | Bd |  |  | Rv |
|  |  |  | Bd |  | Rv |
|  |  |  |  | Bd | Rv |
|  |  |  |  |  | Bd & Rv |
| None |  |  |  |  | Control |

### Timing of parasite exposure

For the six co-infection treatments, timing of the first parasite exposure relative to the second parasite was based on the generation time of the first parasite (Table 1). *A. hamatospicula* takes approximately 11 days to reach the GI tract after initial infection, about 19 days to acquire antibodies, and 24 days to produce offspring (Knutie et al. 2017). Therefore, frogs were first exposed to *hamatospicula* at 24, 19, 11, 1, or 0 days prior to a second parasite exposure. Bd has a generation time of ∼4 days at room temperature, with growth leveling off over time (Piotrowski et al. 2004). Based on our experience with Bd infection in Cuban treefrogs, Bd-induced mortality generally begins about three weeks after exposure. Therefore, to avoid substantial mortality, frogs were exposed to Bd at 16, 12, 8, 4, or 0 days prior to a second parasite exposure. Finally, Ranavirus replicates rapidly, causing mortality in as few as two days for tadpoles and seven days for adults from highly susceptible species, but can also be cleared by immune responses mounted in one to two weeks in other frog species (Green et al. 2002, Gantress et al. 2003, Maniero et al. 2006, Sutton et al. 2014). Consequently, frogs were first exposed to Ranavirus 12, 8, 4, 2, or 0 days prior to a second parasite exposure. Time points were chosen to simulate natural points in disease progression for each parasite (Gray et al. 2009, Voyles et al. 2009, Knutie et al. 2017). Longer time lags between parasites were chosen so that the second parasite exposure occurred when the host was recovering from or in the established stages of infection. Shorter time lags for each parasite were chosen so that the second parasite exposure occurred when the host was in the beginning stages of combating the initial parasite.

### Parasite exposures

Depending on treatment, frogs were exposed in Petri dishes (25 x 100 mm) to a 1-ml aliquot of DI water containing either 30 J3 *A. hamatospicula* larvae, a 1-ml aliquot of DI water containing 10^5^ Bd zoospores, and/or orally to a 77-µl aliquot of MEM containing 10^5^ plaque forming units (PFU) of Ranavirus (Hoverman et al. 2010). Exposures with these doses have been shown to induce sublethal effects in amphibians (Hoverman et al. 2010, Gervasi et al. 2013, Knutie et al. 2017). Sham exposures consisted of the same volume of liquid without the infectious agent for all parasites (1-ml DI water for *A. hamatospicula* and Bd and/or 77-µl of MEM for Ranavirus). All hosts received sham exposures for parasites they were not exposed to. For example, a host singly infected with Bd received sham exposures for Ranavirus and *A. hamatospicula*. Control hosts received sham treatments for all three parasites on day zero. Animals were maintained in the inoculum for 24 h to allow *A. hamatospicula* larvae time to penetrate the host skin (Knutie et al. 2017).

### Amphibian responses to parasites

To measure the growth of amphibian hosts over the duration of the experiment, 24 h after secondary parasite exposure, treefrogs were weighed and snout-vent length (SVL) was measured, and these measurements were taken weekly for four weeks. Survival and disease signs were assessed twice daily. At the time of death, we weighed deceased individuals, measured SVL, and assessed infection for the secondary parasite to which the individual was exposed. Disease signs were determined to be internal hemorrhaging or erythema near the feet for Ranavirus and discoloration, sloughing, or thickening (the latter of which was only noted during dissections) of the skin for Bd (Cunningham et al. 1996, Kilpatrick et al. 2010). For Bd- and Ranavirus-exposed individuals, we swabbed the skin and conducted qPCR (see below). For *A. hamatospicula* exposed individuals, we conducted dissections of the gut. Twenty-eight days after exposure to the secondary parasite, all surviving frogs were euthanized and dissected. Host growth rate was calculated as the final mas or snout-vent length (SVL) minus initial mass or SVL divided by weeks alive (g wk^-1^).

### Assessing parasite loads

To assess *A. hamatospicula* infection, treefrog feces were checked weekly for larval worms and amphibian hosts were dissected at time of morality and at the end of the experiment to quantify adult *A. hamatospicula* infection load in the gastrointestinal tract. To assess Bd and/or viral load, amphibians were swabbed 2, 4, 8, 16, and 24 days after exposure to the second parasite and at the time of mortality. Individuals were swabbed 5 times from hip to toe on both rear legs for Bd and around the cloaca and mouth to test for Ranaviral infection. Swabs were placed in 2-ml sterile microcentrifuge tubes and stored at -80 °C until processing.

DNA was extracted (Qiagen DNEasy Blood & Tissue Kit) from 2-, 4, 8, and 24-day swabs where Ranavirus was the second infecting parasite and from day 4-, 8-, 16-, and 24-day swabs where Bd was the second infecting parasite. Day 2 swabs were not extracted for Bd because Bd DNA from the original inoculum can be detected for up to 2 days after dosing, making it impossible to differentiate infection from exposure at this time (McMahon et al. 2014). Bd and Ranavirus swabs were analyzed using quantitative polymerase chain reaction (qPCR) to quantify parasite load (Boyle et al. 2004, Picco et al. 2007).

### Assessing amphibian immune response

We conducted a separate experiment that included 48 additional Cuban tree frogs to quantify amphibian antibody responses to single and simultaneous co-infections with the parasites that we tested above. Four individuals were tested for each of the three single infection treatments, for a total of twelve singly infected individuals. Eight individuals were tested for each of the three, pairwise, co-infection treatments and half of the individuals in each pairwise co-infection treatment were tested for one parasite and the other half were tested for the other. Finally, twelve individuals were tested as controls, with four individuals tested to compare with each parasite (Table S3). To measure the immune response for each parasite, frogs were euthanized as described above and blood was extracted from frogs with microcapillary tubes 11, 19, and 28 days after exposure to Ranavirus, *A. hamatospicula*, and Bd, respectively. These times were chosen to reflect when antibody immune responses to each parasite should be detectable (Gantress et al. 2003, Ramsey et al. 2010, Knutie et al. 2017). While we acknowledge that immunity is complex, multifactorial response, we chose to focus on IgY antibodies because IgY is a metric of acquired immunity, is the most common immunoglobulin, and is used to combat the parasites we tested (Gantress et al. 2003, Maniero et al. 2006, Knutie et al. 2017, Grogan et al. 2018). General IgY measurements cannot differentiate between Th1 and Th2 immune responses (Bretscher 2014, Menon et al. 2018). However, assuming co-infected individuals would mount higher IgY levels than singly infected individuals to combat both pathogens, no significant increase in IgY levels for co-infected individuals could suggest that a Th1/Th2 tradeoff is limiting the hosts’ ability to mount a significantly higher immune response.

Blood plasma was used in enzyme-linked immunosorbent assays (ELISA) to detect IgY antibody presence. Individual frog serum was first diluted at 1:100 in carbonate coating buffer (0.05 M, pH 9.60) and then added to ninety-six well plates with 100 μL/well. Each individual sample was run in triplicate. After serum was added, plates were incubated at 4 °C overnight. Plates were washed five times with 300 μL/well of a Tris-buffered saline wash solution. The plates were then filled with 200 μL/well of bovine serum albumin (BSA) blocking buffer and incubated for 2 h at room temperature on an orbital table. Plates were washed again five times, loaded with 100 μL/well of primary detection antibody (Goat-αAlligator-IgY, diluted 1:1000; Bethyl), placed on an orbital table for 1 h, washed again (five times), loaded with 100 μL/well of secondary detection antibody (Rabbit-αGoat-IgG, diluted 1:5000; Bethyl) for 1 h, and then washed for the final time (five times). Plates were then loaded with 100 μL/well of tetramethylbenzidine (TMB: Bethyl Laboratories) and incubated for 30 mins before the reaction was stopped with 100 μL/well of stop solution (Bethyl Laboratories). Finally, optical density (OD) of the wells was measured with a spectrophotometer (BioTek, PowerWave HT, 450-nanometer filter).

### Statistical Analyses

All analyses were run with R version 3.4.2 (R Core Team 2017). Plots were created using the *visreg* package and *visreg* function (Breheny and Burchett 2019).

#### Goal 1: Do prior infections and the timing of those infections change the abundance of the secondary parasite?

Importantly, there are two time variables used for Bd and Ranavirus in the statistical analysis. First, loads of Bd and Ranavirus were measured over time after co-infection or single infection. This measurement is not present for *A. hamatospicula* because loads of *A. hamatospicula* were only quantified through destructive sampling at the end of the experiment. Second, there is the time lag between the first and second infection in the co-infection situations. This measurement occurred for all parasites tested.

To test how the interaction between initial parasite identity and time since co-exposure (the amount of time after co-infection) changed parasite growth, we compared the load of the secondary parasite to load of the same parasite species in a single infection treatment through time. These analyses were only done for Bd and Ranavirus because *A. hamatospicula* load data was only collected, at the end of the experiment. We conducted a general linear mixed model (lmer) with the load of the secondary parasite as the dependent variable, initial parasite identity (none if single infection) and swabbing time points (time since co-exposure) as interacting independent variables, and individual as a random effect to account for repeated sampling of the same individuals through time. For Bd, the error distribution was negative binomial because the counts were overdispersed with no upper bound. Viral loads were also counts with no upper bound, but the range of viral loads were too large for the negative binomial model to converge; so, we were forced to use a log-normal model. Tukey’s post-hoc tests were run to compare parasite treatments of co-infecting parasites to one another and to single infections (eg. Bd only, Ranavirus x Bd, *A. hamatospicula* x Bd). All Tukey’s post-hoc tests were run using the *multcomp* package and *glht* function (Hothorn 2010).

To test the effects of the identity of the initial parasite and time lags between parasite exposures (*i.e*. the time gap between co-infecting parasites) on secondary parasite loads (loads measured on day 8 for Bd, day 2 for Ranavirus, day 24 for *A. hamatospicula*), we conducted a generalized linear model (glm) with secondary parasite loads (Bd or Ranavirus) on the first day the secondary parasite load was measured as the negative binomially distributed dependent variable and the identity of the initial parasite and time since initial parasite exposure as interacting independent variables. Zero-inflated negative binomial and Poisson error distributions were also compared, but the negative binomial error distribution was found to have the lowest AIC. For *A. hamatospicula*, we used a logistic regression (*glm* function) with the number of larvae (second infection) that successfully infected (penetrated the host, reached the gut, and reached maturity), out of the total number of larvae to which hosts were exposed, as the dependent variable. Time since initial parasite exposure and the identity of the initial parasite were crossed independent variables. Time since prior exposure in co-infection treatments was a proportion (values between 0 and 1). To calculate this proportion time since exposure for a given parasite treatment was divided by the maximum time since initial exposure possible for each of the parasites (*e.g*. 24, 16, and 12 for *A. hamatospicula*, Bd, and Ranavirus respectively). This allows for comparisons among the various parasites even though the timing of initial exposures were different. We also tested how timing of prior exposure altered parasite growth rate (see supplemental methods and results).

#### Goal 2: Do prior infections and the timing of those infections affect host growth, survival, and tolerance?

To test how the timing of initial infections alters host growth rate, we conducted a general linear model with amphibian growth rate (mass and SVL) over four weeks ([mass_final_ - mass_initial_] /4 weeks) as the dependent variable. Time since initial parasite exposure, the identity of the initial parasite, and the identity of the second (co-infecting) parasite were interacting independent variables and starting weight was a covariate to control for the different host sizes at the start of the experiment. Tukey post-hoc tests were run to compare initial parasite identities to one another and second parasite identities to one another. Dunnett’s tests, using the *multcomp* package and *glht* function (Hothorn 2010), were used to examine if the time lags between the parasite exposures matter when comparing each co-infection time lag to the single infection.

To investigate how the timing of initial infections alters host survival, we conducted a survival analysis using the *survival* package and the *coxph* function (Therneau and Lumley 2019). Individuals that survived to the end of the experiment were right-censored. We included the time since initial exposure, the identity of the initial parasite, and the identity of the second (co-infecting) parasite as interacting independent variables.

To test how the timing of the initial infection and the identity of the co-infecting parasite alters amphibian tolerance (maintaining survival or growth with high parasite loads), we re-conducted the previously described growth and survival analyses with secondary (co-infecting) parasite load (loads measured at the 1^st^ time point; log transformed for Bd or Ranavirus, not log transformed for *A. hamatospicula*) as an additional crossed independent variable.

#### Goal 3: Do co-infections affect host immunity?

To test how co-infections vs. single infections affects host immunity, we conducted a general linear model with antibody levels as the dependent variable, and single vs. co-infection treatments as the independent variable. To test how treatment group (*e.g*. controls, Bd, Bd and Ranavirus, etc.) affects host immunity, we conducted a general linear model with antibody levels as the dependent variable and treatment group as the independent variable. Given that Bd is known to suppress lymphocytes, but not some other immune metrics (Fites et al. 2014), we further tested how Bd impacts host immunity by testing how exposure to Bd alters antibody levels compared to parasites that are not known to be immunosuppressive. We conducted a general linear model (lm) with Bd exposure (*e.g*. bd exposed vs. exposed to other parasites vs. controls) as the independent variable and antibody levels as the dependent variable. Mass of individuals and timing of blood removal were initially used as covariates for all the above analyses, but were excluded in final analyses, as they were not significant. As a reminder, in this portion of the experiment there were no time lags between infections and therefore no analysis of time lags was included.

## Results

### Goal 1: Do prior infections and the timing of those infections change the abundance of the secondary parasite?

Importantly, there are two time variables for Bd and Ranavirus that we cover in this Results section. There is a measurement of time lag between the initial and secondary infection and there is the change in loads of Bd and Ranavirus through time after exposures for both singly and co-infected hosts.

### Ranavirus as the secondary parasite and response variable

We did not find evidence for priority effects when Ranavirus was the secondary parasite, as time lags between parasite exposures did not affect viral loads (Time lag between co-infections: F_1,44_=1.738, *p*=0.194). However, we did find effects of co-infections on the change in viral loads through time. More specifically, we found that the identity of the first infecting parasite (χ^2^_2_=17.2, *p*<0.001), time since hosts were co-infected (χ^2^_2_= 77.65, *p*<0.001), and an interaction between these two factors (Treatment x day: χ^2^_4_ = 12.12, *p*=0.017) affected viral load. Previous infection with *A. hamatospicula* or Bd increased host viral loads relative to loads in hosts exposed to Ranavirus only (Ranavirus vs. Bd and Ranavirus: *p=*0.003; Ranavirus vs. *A. hamatospicula* and Ranavirus: *p*=0.034) and this response was independent of the time since co-exposure for *A. hamatospicula* but increased with time since co-exposure for Bd (Parasite identity: χ^2^_2_=17.2, *p*<0.001; time: χ^2^_2_= 77.65, *p*<0.001; parasite identity x time: χ^2^_4_ = 12.12, *p*=0.017; Fig. 1). However, twenty-four days after co-infection there was no significant difference between viral loads of co-infected and singly infected hosts (*p*=0.23), because the co-infected hosts seemed to begin clearing Ranavirus (Fig. S3B).

**Fig. 1.**
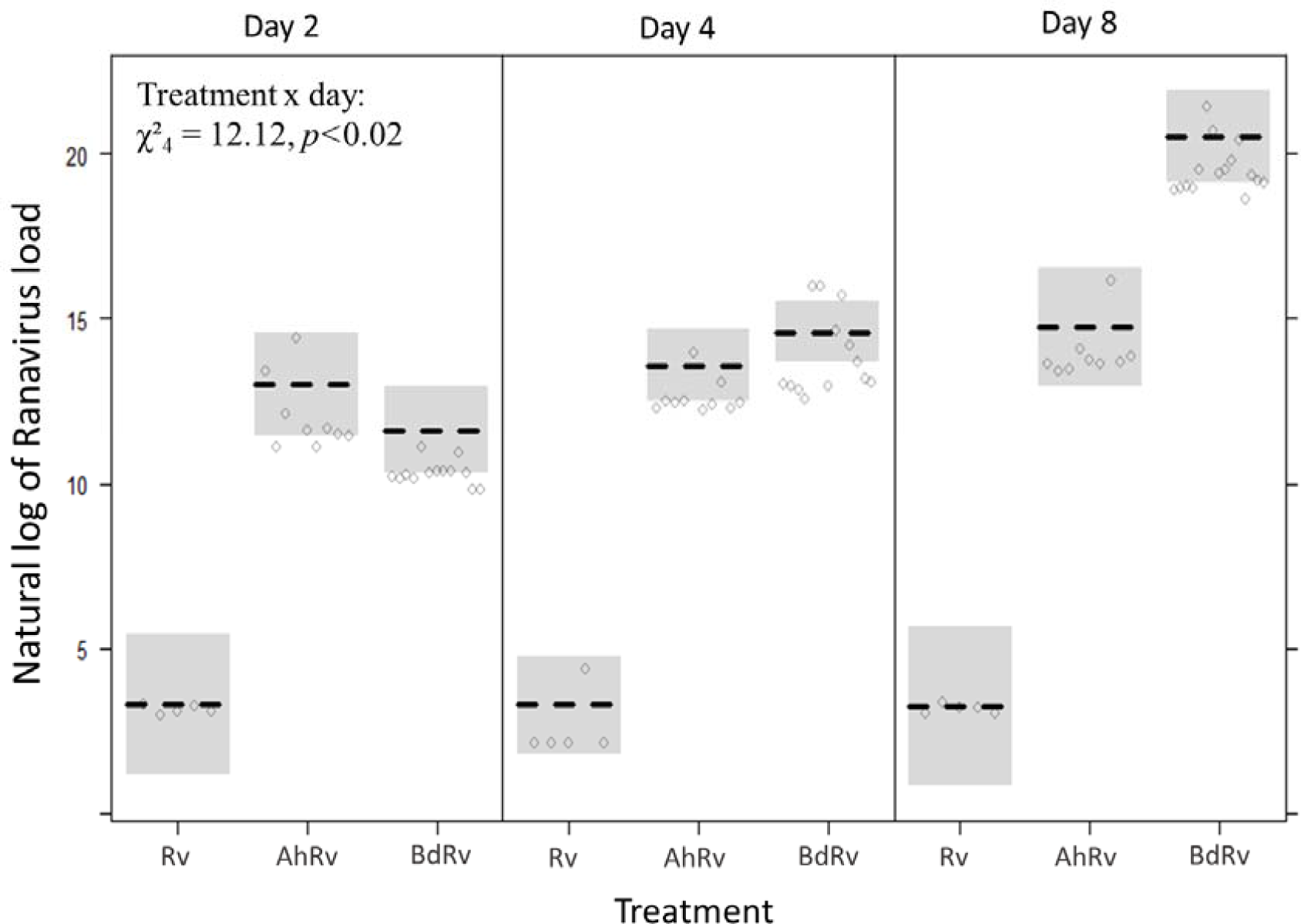
Log-transformed viral load at 2, 4, and 8 days after Ranavirus exposure for *O. septentrionalis* singly infected with Ranavirus (Rv), or initially exposed to *Aplectana hamatospicula* (AhRv) or *Bactrachochytrium dendrobatidis* (BdRv) before Ranavirus exposure. Data are shown as a conditional plot (*i.e*. controlling for everything else in the model), with the expected value (black hashed line) and partial residuals (points) displayed. Days since infection was modeled as a continuous variable. Co-infected hosts had higher viral loads than singly infected hosts (Treatment: χ^2^_4_ =17.24, *p*=0.0002) and these differences increased through time (Treatment x day: χ^2^_4_ = 12.12, *p*<0.02).

### Aplectana hamatospicula *as the secondary parasite and response*

We did not find evidence for priority effects when *A. hamatospicula* was the secondary parasite, as time lags between parasite exposures did not affect *A. hamatospicula* loads (Time lag: χ^*2*^_1_:0.073, *p*=0.78). However, we did find effects of co-infection on *A. hamatospicula* loads. Initial exposure to Bd, regardless of the time lags between co-exposures, was associated with a higher load of *A. hamatospicula* in hosts compared to hosts initially exposed to Ranavirus (Parasite identity: χ^*2*^_1_=9.563, *p*=0.008; Bd and *A. hamatospicula* vs. Ranavirus and *A. hamatospicula*: *p*=0.037; Fig. 2B). *A. hamatospicula* loads in co-infected hosts were not different from loads of hosts infected with *A. hamatopicula* only (Ranavirus and *A. hamatopicula* vs. *A. hamatopicula*: *p*=0.752; Bd and *A. hamatopicula* vs. *A. hamatopicula*: *p*=0.191; Fig. 2B).

**Fig. 2.**
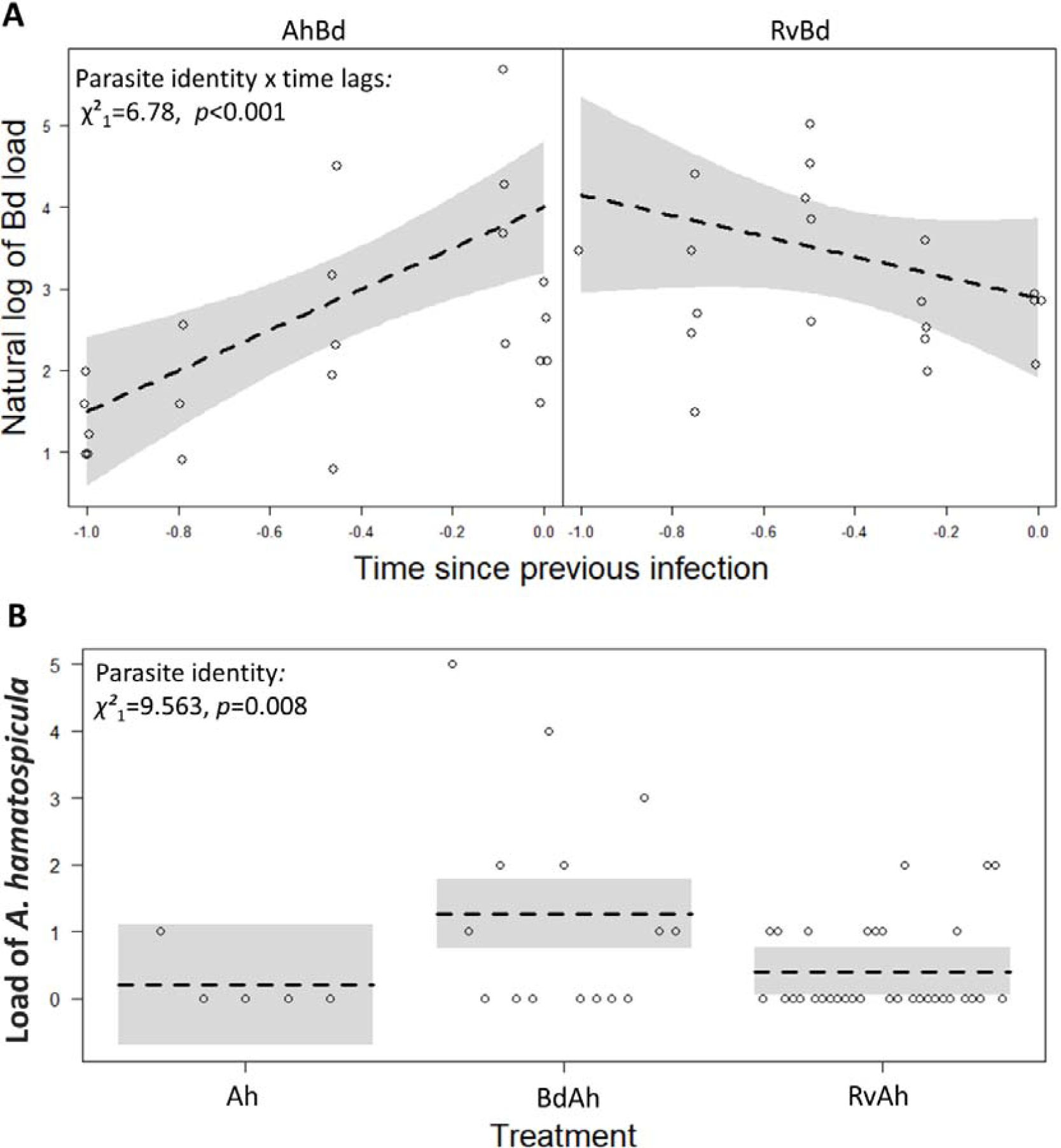
**(A)** *Batrachochytrium dendrobatidis* (Bd) loads on *Osteopilus septentrionalis* hosts (8 days after exposure to Bd) as a function of the standardized time lag (-time lag/maximum time lag) since first exposed to either *Aplectana hamatospicula* (Ah) or Ranavirus (Rv). A -1.0 corresponds to the longest time lag between co-infections and 0 corresponds to simultaneous co-infection. The time lag between co-infecting parasites was associated positively with Bd loads for Ah (*p*=4.30e-5), but negatively for Rv (*p*=0.058). **(B)** Prior exposure to Bd allowed more Ah to successfully establish in hosts than prior exposure to Ranavirus (Initial parasite identity: χ^*2*^_1_=9.563, *p*=0.008), but singly infected hosts did not have significantly different loads from either co-infected host treatments (*p*>0.190). Data in both panels are show as conditional plots (*i.e*. controlling for everything else in the model), with the expected value (black hashed line), a 95% confidence bands or intervals for the expected value (gray band), and partial residuals (points) displayed.

### Batrachochytrium dendrobatidis *as the secondary parasite and response*

There was evidence for priority effects when Bd was the secondary parasite (Parasite identity x time lag: χ^2^_1_:6.78, *p*=0.009). When *A. hamatospicula was* the first parasite, time lags between parasite exposures and Bd loads at 8 d were correlated negatively (*Z*=4.09, *p*<0.001). In contrast, when Ranavirus was the first parasite, time lags between parasite exposures and Bd loads at 8 d were correlated positively (*Z*=-1.89 *p*=0.058; Fig. 2A). When looking at loads of Bd through time since co-infection, loads of Bd were positively associated with time (Swab day: χ^2^_1_=173.78, *p*<0.001), but these trends were not affected by co-infection or the identity of the co-infecting parasite (Swab day x treatment: χ^2^_2_: 1.65, *p*=0.439; Fig. S3A).

### Goal 2: Do prior infections and the timing of those infections affect host growth, survival, and tolerance?

#### Growth as the response

Longer time lags between parasite exposures were associated with lower host growth based on SVL (mm/week) when both Bd and Ranavirus were the initial parasites, regardless of whether these hosts were infected singly or co-infected (*F*_1,49_=5.00, *p*=0.03; *F*_1,45_=6.617, *p*=0.013, respectively; Fig. S1). The growth rates of co-infected (at all initial infection time points) and singly infected hosts did not differ significantly at any time since exposure (Fig. S1; Table S1). These effects were similar when growth rates were calculated based on weight (see supplemental results).

#### Survival as the response

Time lags between parasite exposures did not affect the survival of hosts (χ^2^_1_=1.72, *p=* 0.19). However, when hosts were infected initially with Bd or Ranavirus, hosts had a significantly lower survival rate than hosts initially infected with *A. hamatospicula* (Identity of first parasite: χ^2^_2_=22.92, *p*<0.001; Bd vs. *A. hamatospicula*: *p*=0.003; *Ranavirus vs. A. hamatospicula: p*=0.043; Figure 5). Bd, Ranavirus, and *A. hamatospicula* did not significantly affect host survival if they were the second infecting parasite (Identity of the second parasite: χ^2^_2_=2.85, *p=*0.24).

#### Tolerance as a response

Time lags between parasite exposures did not alter host tolerance (the slope between load and growth rate [g/week] or survival) to secondary infections (Viral load x time lag: Host Growth: *F*_1,35_=0.19, *p*=0.665; Host survival: χ^2^_1_=1.18, *p=*0.28; see supplement for tolerance to Bd and *A. hamatospicula*). However, the identity of the initial parasite did impact host tolerance to the secondary parasite. Previous infection with *A. hamatospicula* reduced host tolerance (lower growth rate [g/week] and lower survival at higher loads) of Ranavirus infections more than did previous Bd infections (Viral load x identity of second infecting parasite: Host growth: *F*_1,35_= 6.033, *p*<0.02; Fig. 3; Host survival: χ^2^_1_=6.98, *p=*0.008). In fact, hosts that were exposed to Bd and then Ranavirus were hyper-tolerant of Ranavirus as growth rates were positively correlated with viral loads (Fig. 3). Despite this, hosts infected with Bd before Ranavirus were not significantly more tolerant of Ranavirus infections than hosts singly infected with Ranavirus (Co-infection with Bd x viral load: *F*_1,22_=0.4, *p*=0.8). Co-infection and timing of co-infections did not change host tolerance to Bd, *A. hamatospicula* (host growth or survival), or Ranavirus (survival; see table S2).

**Fig. 3.**
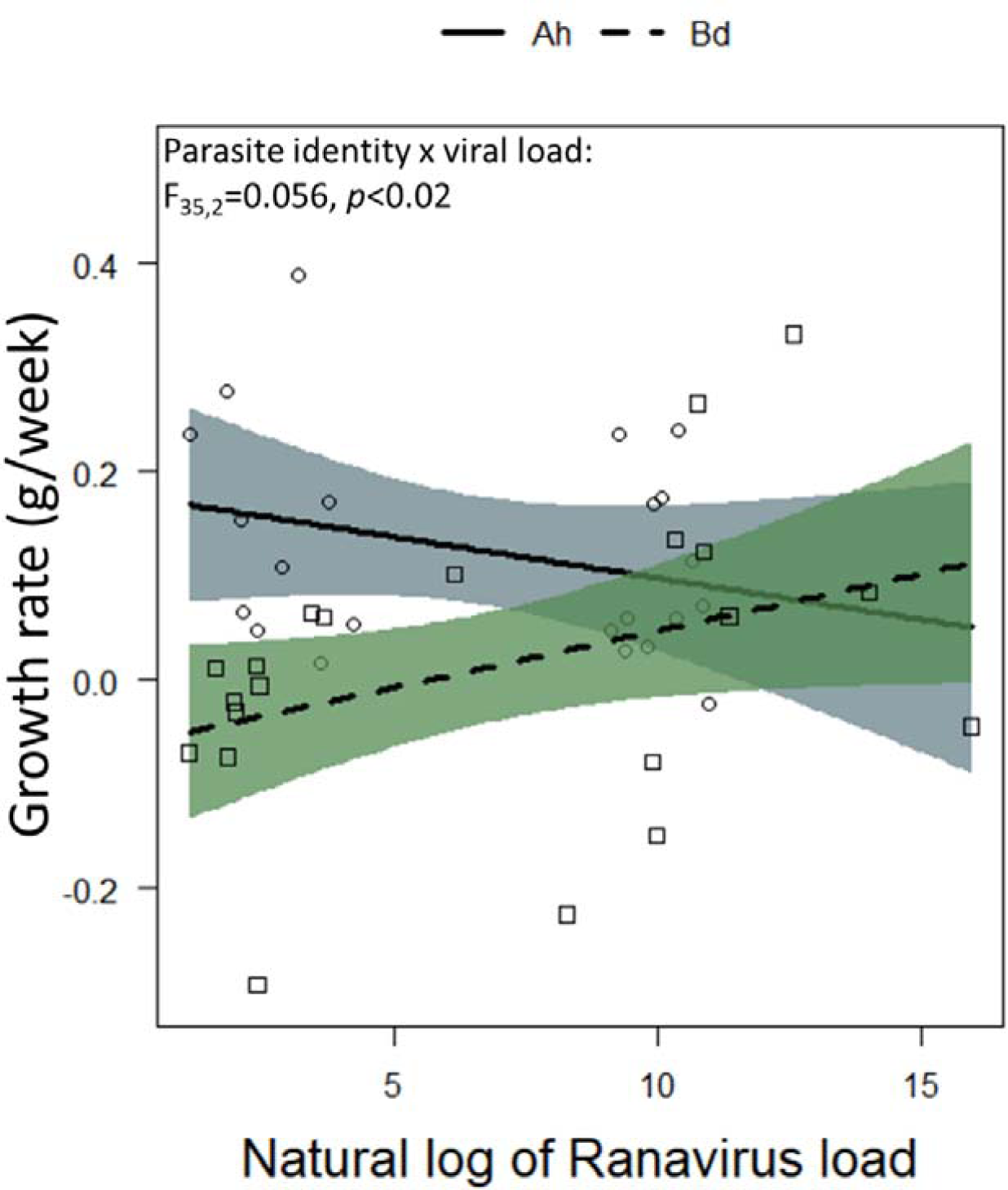
*Osteopilus septentrionalis* tolerance (growth rate as a function of viral load) of loads of Ranavirus when first exposed to either *Aplectana hamatospicula* (Ah) or *Batrachochytrium dendrobatidis* (Bd). Data are shown as a conditional plot (*i.e*. controlling for everything else in the model), with the expected value (the black line and black dotted line represent *A. hamatospicula* and Bd respectively), a 95% confidence band for the expected value (light and darker grey shading represent *A. hamatospicula* and Bd respectively), and partial residuals (points) displayed. Initial infection with Ah or Bd decreased and increased host tolerance, respectively (Initial parasite identity x viral load: F_35,1_=0.056, *p*<0.02).

#### Signs of disease

Signs of Bd (*i.e*., skin thickening, sloughing, or discoloration) was most common when Bd infected the host first and therefore had more time to reach higher loads. Signs of Ranavirus infection (*i.e*., hemorrhaging) were present in all co-infections, regardless of whether Ranavirus infected first or second. Nineteen of the 35 frogs that experienced mortality showed signs of disease, 9 individuals showed signs of Ranavirus, 5 of Bd, and 5 of both parasites (Table S4).

### Goal 3: Do co-infections affect host immunity?

The antibody response of hosts was not altered by co-infections when compared to single infections or controls (Number of infections: *F*_2,39_=1.98, *p*=0.152) and did not differ significantly when comparing all treatment groups (Treatment group: *F*_6,35_=1.43, *p*=0.232; Fig. S2). Bd is documented to suppress acquired immune function (Rosenblum et al. 2009, Fites et al. 2013, Fites et al. 2014), whereas the other two parasites are not. Thus, we hypothesized that Ranavirus and *A. hamatospicula* would elevate antibodies relative to controls, but that Bd might not. This hypothesis was supported as exposure to Ranavirus and *A. hamatospicula* (grouping individuals co-infected and singly infected with each parasite) increased antibody levels relative to control animals (Parasite identity: *F*_2,39_=4.15, *p*=0.024; Ranavirus and *A. hamatospicula* vs. control: *p*=0.024), but Bd did not (Bd vs. control: *p*=0.709; Fig. 5). We did not stagger time lags between co-infections in this portion of the experiment and therefore found no evidence for or against priority effects altering antibody levels.

## Discussion

Organisms in the wild are regularly co-infected with several species of parasites. Furthermore, these co-infections are likely to be staggered and thus it is important to understand if the time lags between these parasite exposures affect disease progression and host health, a primary goal of this work. Our study found that the identity of the co-infecting parasite, and the time lag between co-infecting parasites can alter disease progression in hosts (Table 2).

**Table 2.**
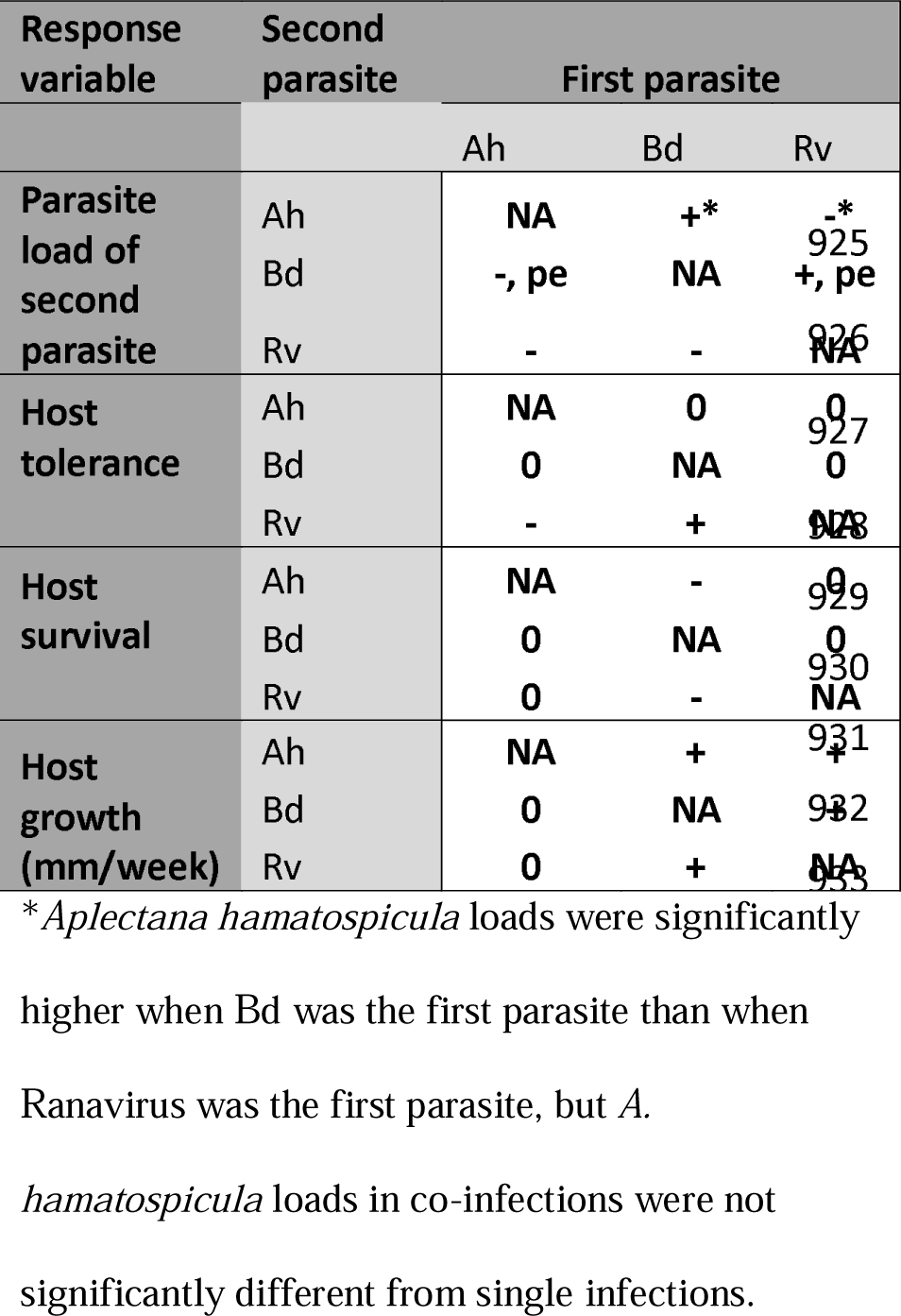
A summary of the effects of parasite identity and time lag between co-infecting parasites on the abundance of the second parasite and host tolerance of infection, survival, and growth. Ah = *Aplectana hamatospicula*, Bd = *Batrachochytrium dendrobatidis*, Rv = Ranavirus, + = significant positive effect, - = significant negative effect, 0 = non-significant effect, pe = significant priority effect (significant association between the time lag between co-infecting parasites and the load of the second parasite)

Hosts co-infected with *A. hamatospicula* and Ranavirus, had increased viral loads and decreased *A. hamatospicula* loads relative to single infections, regardless of the timing of those infections (Fig. 1,2). Therefore, for this co-infection pairing, we found evidence of parasite interactions, but not priority effects. One of the potential drivers for these interactions could be in-direct interactions mediated by the host immune system. In many vertebrate species, Th1 and Th2 immune responses suppress one another and thus hosts have difficulty mounting simultaneous defenses against macro and microparasites (Abbas et al. 1996, Morel and Oriss 1998, Yazdanbakhsh et al. 2002). Indeed, helminth infections can suppress the INF-γ expression that promotes the Th1 arm of the immune system, which targets most intracellular microparasites (Graham 2008). Consistent with the notion that Th1 and Th2 are mutually exclusive and thus tradeoff was the lack of an additive IgY antibody response in hosts co-infected with Ranavirus and *A. hamatospicula* when compared to single infections with these parasites (Fig. S4). Therefore, higher Ranavirus infection loads following *A. hamatospicula* exposure may be driven by the host having previously mounted an immune response that not only limits available resources to combat a second parasite, but also suppresses the immune response necessary to combat Ranavirus. While this is one potential mechanism for this pattern, IgY is not the only way hosts combat parasites. Other aspects of immunity, such as antimicrobial peptides limiting the ability of parasites to enter the skin, or non immune mechanisms, such as amphibians eating more to lessen the negative effects of infections, might have also played a role in this outcome (Rollins-Smith et al. 2002, Hess et al. 2015, Knutie et al. 2017). Especially since hosts are clearing viral loads at day twenty-four and therefore appear to eventually be able to mount an effective acquired immune response (Fig. S3B). Conversely, *A. hamatospicula* loads were lower with Ranavirus infection. This result could be mediated by direct competition between parasites. Ranavirus infections commonly induce hemorrhaging of intestinal epithelial cells, potentially compromising the resources or living space of *A. hamatospicula* (Miller et al. 2011). If this direct effect was stronger than the indirect effect of a tradeoff between the Th1 and Th2 immunity, then the effect of Ranavirus on *A. hamatospicula* would be inhibitory rather than facilitative. These results suggest that macroparasite co-infections could increase Ranavirus loads (except see Wuerthner et al. 2017), which could significantly impact amphibian population health (Green et al. 2002).

In co-infections with Bd and *A. hamatospicula, A. hamatospicula* loads increased and Bd loads decreased when they were the second parasite (Fig.2). *A. hamatospicula* enters through the skin of the frog (Knutie et al. 2017), which is also the site of Bd infection (Berger et al. 2005). Therefore, higher *A. hamatospicula* loads after Bd infection suggests that Bd infection facilitates the process of juvenile worms entering the skin. Several mechanisms are possible for this facilitation. First, Bd disrupts the normal functioning of the skin (Voyles et al. 2009, Kilpatrick et al. 2010), and can be locally immunosuppressive of lymphocyte proliferation (Fig. 5; Fites et al. 2013). Therefore, *A. hamatospicula* loads could be higher after Bd exposure due to direct interactions between these parasites or indirect interactions from Bd immune suppression. When the order of infections was reversed, Bd loads were lower the longer the host was previously infected with *A. hamatospicula* (Fig. 2), a priority effect. Given that *A. hamatospicula* establishes as adults in the gut (Knutie et al. 2017) but Bd infects the skin (Berger et al. 2005), this priority effect is unlikely to be a product of direct interactions between these parasites once *A. hamatospicula* establishes. We predicted that Bd loads would be facilitated by previous *A. hamatospicula* infection because of the Th1/Th2 immune tradeoff. Although fungal infections cause the host to upregulate the Th1 arm of the immune system (Blanco and Garcia 2008), they also frequently, but not always (Zelante et al. 2009), upregulate the Th17 arm of the immune system. Whether amphibians uses the Th1 arm, the Th17 arm, or a mix of the two to combat Bd infections has been debated and is still unknown because of a limited understanding of whether the Th17 arm of the immune system is present in amphibians (Fites 2014). Additionally, other aspects of immunity, such as increased nutrient intake to limit the negative effects of infections or skin sloughing to attempt to reduce Bd loads, could be playing a role in this outcome (Cramp et al. 2014, Hess et al. 2015, Knutie et al. 2017). Consequently, amphibian immune responses, especially those combatting Bd, will need to be better understood to elucidate any immune mechanisms that might have inhibited Bd growth after *A. hamatospicula* infections. If the pattern seen here with *A. hamatospicula* and Bd holds up with other macroparasite infections, macroparasite co-infections (when macroparasites infect first) could have significant positive impacts on lowering the total Bd load in an environment, positively impacting frog populations in the field (Kilpatrick et al. 2010).

In co-infections with Ranavirus and Bd, co-infection with either parasite increased the load of the other. Additionally, there was a positive relationship between Bd loads and the time lag between Ranavirus and Bd infections; the timing of the prior Ranavirus infection altered Bd loads (*i.e*., a priority effect, Fig. 2). In contrast, viral loads increased regardless of the timing of the Bd co-infection (Fig. 1), until hosts had time to mount an acquired immune response to Ranavirus (Fig. S3B). These two parasites are unlikely to interact directly, as Bd infects the skin (Berger et al. 2005) and Ranavirus infects internal organs (Cunningham et al. 1996, Gantress et al. 2003). However, they have the potential for indirect interactions. Infection with Bd significantly suppresses metrics of the hosts’ acquired immune response (Fig. 5; Fites et al. 2013) and this could facilitate subsequent viral infection. If Bd activates the immune system through the Th17 arm of the immune system (Zelante et al. 2009) in addition to the Th1 arm (Blanco and Garcia 2008), then tradeoffs and complexities of mounting these immune responses could account for the facilitative pattern observed in this co-infection pairing. Additionally, these parasites replicate internally and can be costly and deadly to amphibian hosts (Berger et al. 1998, Green et al. 2002, Gantress et al. 2003, Skerratt et al. 2007, Kilpatrick et al. 2010). Given that immune responses are costly and can be traded off with other demanding processes, such as reproduction and growth (Lochmiller and Deerenberg 2000), mounting an immune response sufficient to combat both of these parasites could have been beyond the resource capacity of our hosts. Field studies have found that Ranavirus and Bd are not associated with each other at the site (Olori et al. 2018, Stutz et al. 2018) or individual level, but there are positive associations between these parasites at the species level for some of the sampled amphibian species (Stutz et al. 2018). Despite this, co-infections with Bd and Ranavirus, are common in nature (Whitfield et al. 2013, Warne et al. 2016, Watters et al. 2018) and our findings suggest that these co-infections can be facilitative and thus appear to put host populations at greater risk than single infections.

When we tested how the time lags between infections and the identity of the co-infecting parasites altered host growth, survival, and tolerance, we found that infection with Bd or Ranavirus as the first parasite caused higher mortality than *A. hamastopicula* infections (Fig. 4). We also found that the longer hosts were infected with Bd and Ranavirus, the lower the growth rate of hosts (Fig. S2). These results support our hypothesis that host growth and survival should be lower if the infecting parasite is virulent, like Bd and Ranavirus (Chinchar 2002, Voyles et al. 2009), and has longer to establish. Additionally, hosts were less tolerant of Ranavirus infections (lower growth rate [g/week] with higher viral loads) when the previous infecting parasite was *A. hamatospicula* but were more tolerant when hosts were exposed to Bd before Ranavirus (Fig.4). Hosts not only tolerated viral loads when co-exposed to Bd, but also grew more rapidly at higher viral loads than at lower viral loads when co-infected by Bd. Amphibians display increased growth and development when exposed to other inhospitable conditions such as rapidly drying ponds (Bekhet et al. 2014), pesticides (Rohr et al. 2004), or predators (Relyea 2007), which could mean that hosts are using their resources to grow rapidly when exposed to two deadly amphibian parasites rather than attempting to resist or clear the infections (*i.e*., a life-history trade-off). This is further supported by the fact that this treatment group had the highest proportion of individuals showing signs of disease (Table S4).

**Fig. 4.**
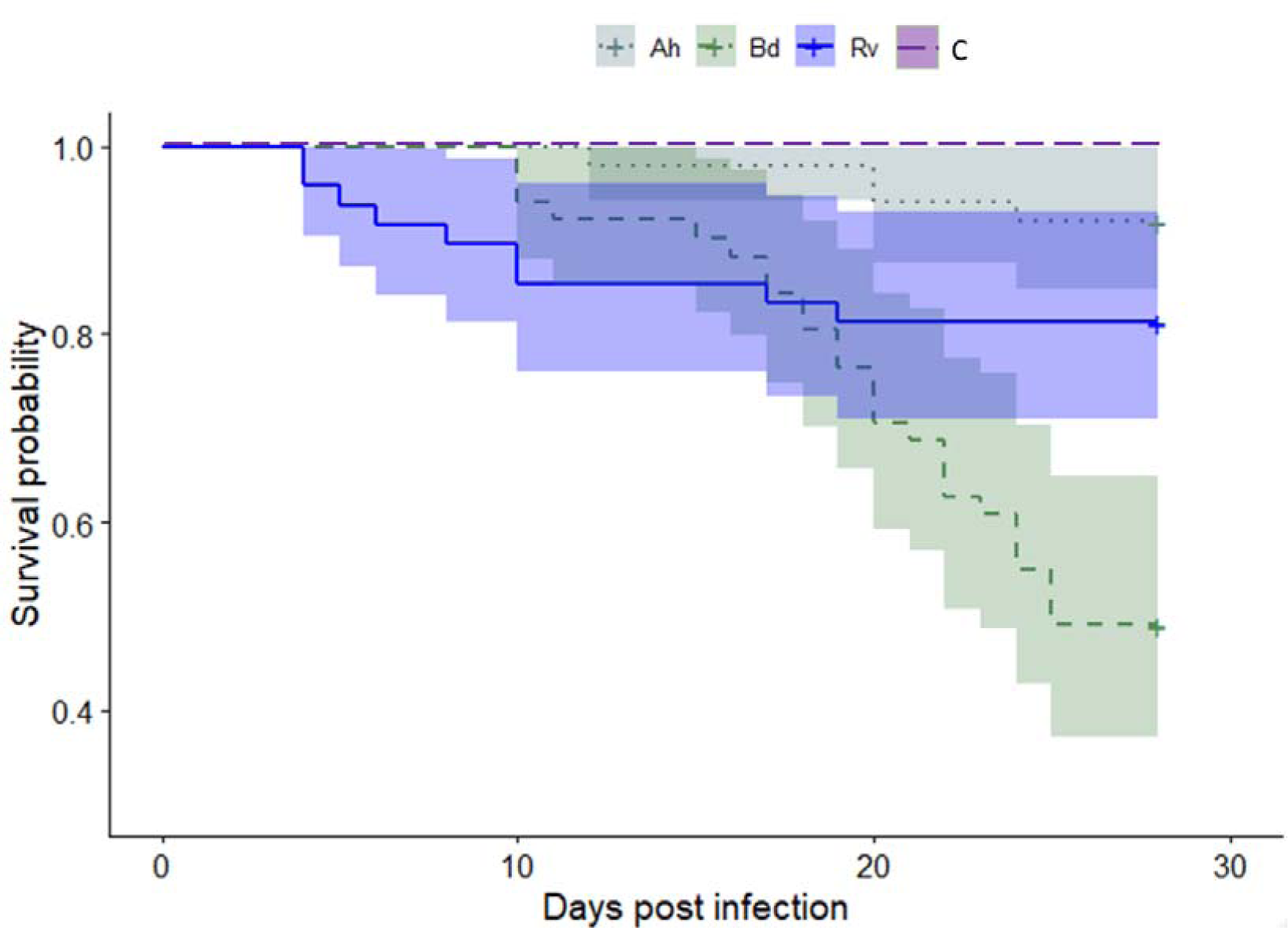
Survival of *Osteopilus septentrionalis* hosts exposed to no parasites (controls, C) or first exposed to *Aplectana hamatospicula* (Ah), *Bactrachochytrium dendrobatidis* (Bd), or Ranavirus (Rv) averaged across both secondary parasites. Data are shown as survival curves, with the median survival (solid and dashed lines) and the 95% confidence interval for the expected value (grey band) displayed. Initial exposure to Bd and Rv reduced days alive relative to initial exposure to Ah (Bd vs. Ah: *p*=0.003; Rv vs. Ah: *p*=0.043).

**Fig. 5.**
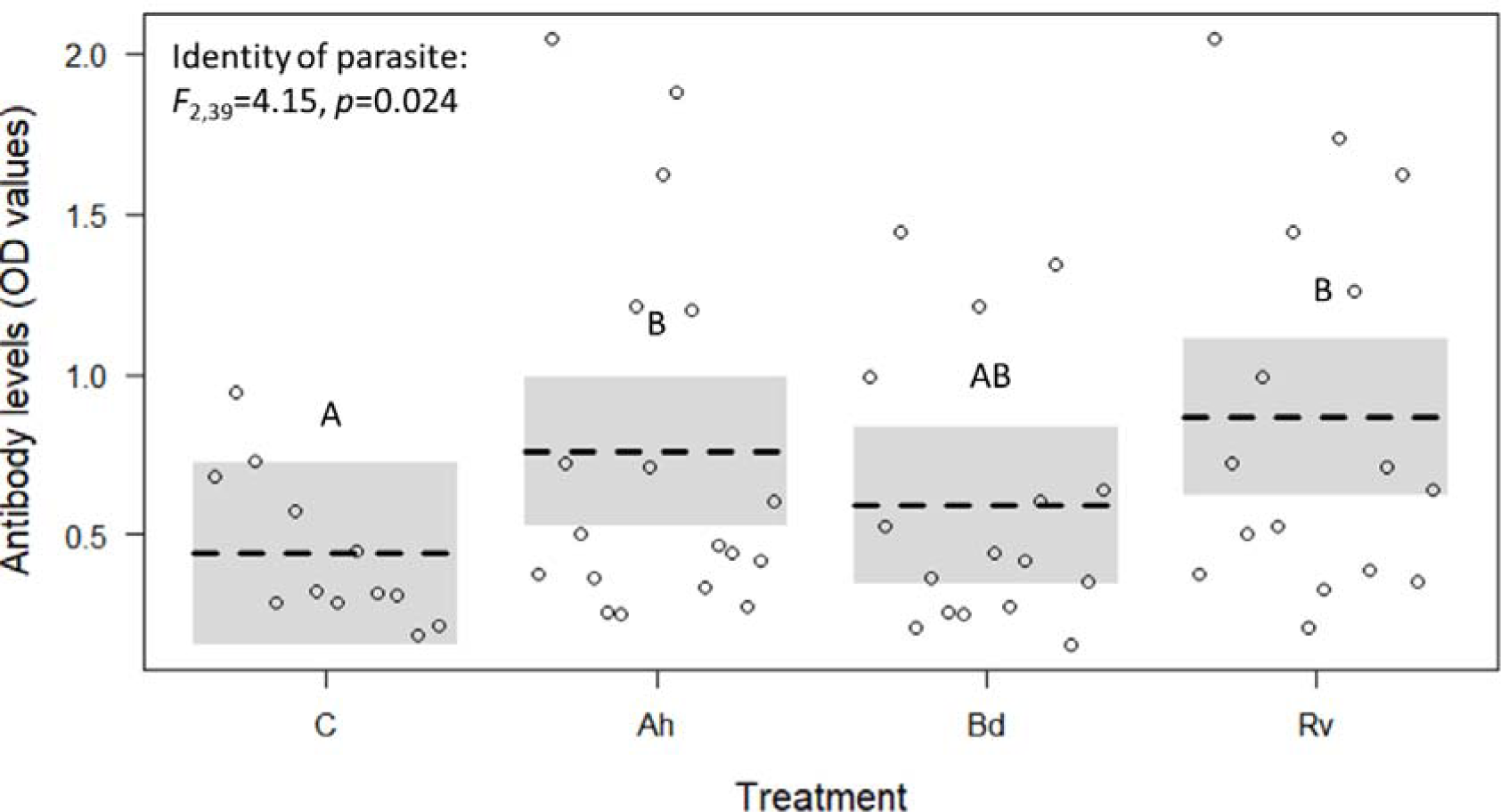
IgY antibody levels for hosts exposed to *Bactrachochytrium dendrobatidis* (Bd), *Aplectana hamatospicula* (Ah), Ranavirus (Rv) averaged across single infections and pairwise co-infections and control individuals with no infections (C). Data are shown as a conditional plot (*i.e*. controlling for everything else in the model), with the expected value (black hashed line), a 95% confidence interval for the expected value (gray band), and partial residuals (points) displayed. Antibody levels were not significantly different in hosts not exposed to parasites and those exposed to Bd (*p*=0.709), which is known to be immunosuppressive (Fites et al. 2014). However, antibody levels were significantly higher in hosts exposed to parasites other than Bd (*i.e*., those not known to be immunosuppressive) relative to controls (Identity of parasite: *F*_2,39_=4.15, *p*=0.024; non-Bd vs. control: *p*=0.024).

Studies broadly support that the order of co-infections can alter parasite communities (de Roode et al. 2005, Devevey et al. 2015, Leopold et al. 2017), but our research revealed that time lags between these co-infections can affect parasite loads, likely as a result of indirect competition mediated by the host immune system. Additionally, host tolerance, growth, and survival can be impacted by these co-infections. Our results suggest that co-infections with Ranavirus and Bd could lead to negative effects on amphibian populations. Also, if our results with *A. hamatospicula* are generalizable to other macroparasite co-infections (see Wuerthner et al. 2017 for an exception), macroparasite co-infections could increase Ranavirus loads and decrease Bd loads. This suggests that co-infections with macroparasites and these microparasites could have a positive or negative overall effect on amphibian population health depending on host susceptibility to these two deadly microparasites. Additionally, while this study was conducted at an intermediate temperature, higher and lower temperatures would likely alter the progression of these diseases (Cohen et al. 2017, Price et al. 2019), potentially changing how time lags between co-infections alter amphibian health. Given this, further research is needed to understand the generality of time lag effects and the mechanisms driving time lag effects on co-infections. A better understanding of when time lags between co-infections are important will guide researchers and practitioners in understanding when co-infections might be particularly problematic. This knowledge may, in turn, allow practitioners to implement more efficient control strategies targeted at co-infections for amphibians and other important hosts such as pollinators (Bromenshenk et al. 2010), livestock (Ezenwa and Jolles 2015), and humans (Corbett et al. 2003).

## Acknowledgements

We would like to thank J. Hoverman for supplying the Ranavirus, S. Knutie for guidance with immune assays, J. Cohen for general guidance, and all the undergraduates who made this experiment possible. Funds were provided by grants to J.R.R. from the National Science Foundation (EF-1241889), the National Institutes of Health (R01GM109499, R01TW010286-01).

