## supplement for "The application of community ecology theory to co-infections in wildlife hosts"

**Methods**

*Statistical Analyses: Do prior infections and the timing of those infections change the growth rate of the secondary parasite?*

To determine how the timing of initial infections alters the growth rates of secondary parasites (Bd and Ranavirus only), we ran a linear model with a Gaussian distribution. We used time since initial parasite exposure and the identity of the initial parasite as crossed independent variables and secondary parasite load growth as the dependent variable. Time since prior exposure in co-infection treatments was a proportion (values between 0 and 1); time since exposure for a given parasite treatment divided by the maximum time since initial exposure possible for each of the parasites (*e.g.* 16, and 12 for Bd, and Ranavirus respectively). This allows for comparisons among the various parasites even though the timing of initial exposures were different. For Ranavirus and Bd, parasite growth rates were quantified by fitting a logistic growth model to mean Bd zoospore counts or Ranavirus viral copies/ng DNA at each swabbing time point (assuming no growth at t 0; bbmle package (Bolker 2017), mle2 function, negative log-likelihood function, assuming a normal error distribution). Tukey post-hoc tests were run to compare how initial parasite identities altered the load of the second parasite.

**Results**

*Goal 1: Do prior infections and the timing of those infections change the growth rate of the secondary parasite?*

Growth rates of Bd were not altered by the timing of the previous infection (*F*_1,39_=0.00, *p=*0.995), the identity of the co-infecting parasite (*F*_1,39_=0.123 *p=*0.73), or their interaction (*F*_1,39_=0.596 *p=*0.21).

Growth rates of Ranavirus were also not altered by the timing of the previous infection (lm: *F*_1,44_=0.05, *p=*0.82), the identity of the co-infecting parasite (*F*_1,44_=0.055, *p=*0.82), of the interaction of the two (*F*_1,44_=1.74, *p=*0.19).

*Goal 2: Do prior infections and the timing of those infections affect host growth and tolerance?*

Host growth rate (g/week) was significantly impacted by the identity of the initial infecting parasite (*F*_2,120_=14.42, *p*<0.001), with an initial infection with Bd suppressing host growth rate in comparison to initial infection with *A. hamatospicula* (*p*=0.004) and Ranavirus (*p*<0.001). Prior infections and the timing of those infections did not impact host ability to tolerate Bd and *A. hamatospicula* infections (see table S2).

**Table S1**. There were no significant differences in growth rate (mm/week) when using Dunnett’s tests (Hothorn et al. 2017) to individually compare each pairwise co-infection at each time point to single infection at time zero. *Aplectana hamatospicula* = Ah, *Bactrachochytrium dendrobatidis*= Bd, Ranavirus = Rv.

| **Ah** as the second parasite | | | |
| --- | --- | --- | --- |
| Co-infection with **Bd** at various time points | | | |
| Coefficient | Estimate | *t* value | *p* value |
| Bd at time 0 | 0.17 | 0.81 | 0.89 |
| Bd at time 0.375 | -0.15 | -0.7 | 0.94 |
| Bd at time 0.7 | -0.05 | -2.23 | 1 |
| Bd at time 0.75 | -0.25 | -1.16 | 0.68 |
| Bd at time 1 | -0.15 | -0.7 | 0.94 |
| Co-infection with **Rv** at various time points | | | |
| Coefficient | Estimate | *t* value | *p* value |
| Rv at time 0 | 0.38 | 1.51 | 0.46 |
| Rv at time 0.25 | -0.25 | -1 | 0.79 |
| Rv at time 0.5 | 0.25 | 1.1 | 0.75 |
| Rv at time 0.75 | -0.25 | -1 | 0.79 |
| Rv at time 1 | -0.15 | -0.64 | 0.96 |
| **Ranavirus** as the second parasite | | |  |
| Co-infection with **Bd** at various time points | | | |
| Coefficient | Estimate | *t* value | *p* value |
| Bd at time 0 | 0.83 | 1.63 | 0.318 |
| Bd at time 0.38 | 0.63 | 1.24 | 0.552 |
| Bd at time 0.75 | 0.03 | 0.08 | 1 |
| Bd at time 1 | 0.03 | 0.07 | 1 |
| Co-infection with **Ah** at various time points | | | |
| Coefficient | Estimate | *t* value | *p* value |
| Ah at time 0 | 2.04E-17 | 0 | 1 |
| Ah at time 0.08 | -0.3 | -1.26 | 0.6 |
| Ah at time 0.46 | -0.2 | -0.84 | 0.87 |
| Ah at time 0.8 | -0.5 | -2.1 | 0.17 |
| Ah at time 1 | -0.3 | -1.26 | 0.6 |
| **Bd** as the second parasite | |  |  |
| Co-infection with **Rv** at various time points | | | |
| Coefficient | Estimate | *t* value | *p* value |
| Rv at time 0 | 0.83 | 1.78 | 0.3 |
| Rv at time 0.25 | 0.13 | 0.29 | 1 |
| Rv at time 0.5 | 0.03 | 0.07 | 1 |
| Rv at time 0.75 | -0.17 | -0.36 | 1 |
| Rv at time 1 | -0.17 | -0.36 | 1 |
| Co-infection with **Ah** at various time points | | | |
| Coefficient | Estimate | *t* value | *p* value |
| Ah at time 0 | 0.23 | 0.73 | 0.93 |
| Ah at time 0.08 | 0.43 | 1.35 | 0.55 |
| Ah at time 0.46 | -0.07 | -0.21 | 1 |
| Ah at time 0.8 | 0.03 | 0.1 | 1 |
| Ah at time 1 | 0.53 | 1.66 | 0.36 |

**Table S2.** A host can tolerate a disease if they can remain healthy when infected with high loads of a parasite. When testing if the timing of the initial parasite infection or the identity of the co-infecting parasite alter a host’s tolerance to *Bactrachochytrium dendrobatidis* (Bd) **(A),** *Aplectana hamatospicula* (Ah) **(B),** or Ranavirus (Rv) **(C)** (mm/week only) we found no effects of either of the interaction of these predictors with parasite load (*eg.* time since initial infection x load of Bd) on host growth (g/week or mm/week) or survival.

**A.**

| **Bd as the second parasite** | | |  |
| --- | --- | --- | --- |
| **Response: Growth Rate (g/week)** | | | |
| Coefficient | *F* value | *p* value | df |
| time since initial infection | 0 | 0.98 | 1,32 |
| identity of initial parasite | 1.4 | 0.25 |  |
| load of Bd | 0 | 0.98 |  |
| load of Bd*time since initial infection | 0.03 | 0.43 |  |
| load of Bd*identity initial parasite | 0.39 | 0.86 |  |
| **Response: Growth Rate (mm/week)** | | | |
| Coefficient | *F* value | *p* value | df |
| time since initial infection | 1.15 | 0.29 | 1,35 |
| identity of initial parasite | 0.18 | 0.67 |  |
| load of Bd | 0.25 | 0.62 |  |
| load of Bd*time since initial infection | 0.86 | 0.36 |  |
| load of Bd*identity of initial parasite | 0.99 | 0.33 |  |
| **Response: Survival** | |  |  |
| Coefficient | χ² value | *p* value | df |
| time since initial infection | 0.73 | 0.39 | 1 |
| identity of co-infecting parasite | 3.06 | 0.08 |  |
| load of Bd | 3.26 | 0.07 |  |
| load of Bd*time since initial infection | 0.24 | 0.62 |  |
| load of Bd*identity of initial parasite | 1.33 | 0.25 |  |

**B.**

| **Ah as the second parasite** | | |  |
| --- | --- | --- | --- |
| **Response: Growth Rate (g/week)** | | | |
| Coefficient | *F* value | *p* value | df |
| time since initial infection | 0.72 | 0.4 | 1,35 |
| identity of co-infecting parasite | 10.11 | 0.003 |  |
| load of Ah | 0.03 | 0.87 |  |
| load of Ah*time since initial infection | 0.22 | 0.64 |  |
| load of Ah *identity of initial parasite | 0 | 0.94 |  |
| **Response: Growth Rate (mm/week)** | | | |
| Coefficient | *F* value | *p* value | df |
| time since initial infection | 1 | 0.32 | 1,38 |
| identity of co-infecting parasite | 0 | 0.95 |  |
| load of Ah | 0.6 | 0.44 |  |
| load of Ah*time since initial infection | 0.67 | 0.42 |  |
| load of Ah*identity of initial parasite | 0.23 | 0.63 |  |
| **Response: Survival** | |  |  |
| Coefficient | χ² value | *p* value | df |
| time since initial infection | 5.28 | 0.02 | 1 |
| identity of co-infecting parasite | 2.63 | 0.11 |  |
| load of Ah | 0.11 | 0.74 |  |
| load of Ah*time since initial infection | 0.08 | 0.78 |  |
| load of Ah*identity of initial parasite | 2.13 | 0.14 |  |

**C.**

| **Rv as the secondary parasite** | | |  |
| --- | --- | --- | --- |
| **Response: Growth Rate (mm/week)** | | | |
| Coefficient | *F* value | *p* value | df |
| time since initial infection | 3.91 | 0.05 | 1,40 |
| identity of co-infecting parasite | 2.29 | 0.14 |  |
| load of Rv | 0.03 | 0.87 |  |
| load of Rv*time since initial infection | 0.35 | 0.56 |  |
| load of Rv*identity of initial parasite | 0.07 | 0.8 |  |

**Table S3**. Hosts were sampled for immune response at time points pertinent to the parasite exposure. Days sampled are shown for all of the below co-infection and single infection treatments and controls. There were 4 individuals for each treatment group shown below. For this purposes of this figure *Bactrachochytrium dendrobatidis* is Bd, Ranavirus is Rv, *Aplectana hamatospicula* is Ah, and controls are C.

|  |  | **Number sampled** | |  |
| --- | --- | --- | --- | --- |
| **Exposures** | d 11 | d 19 | d 28 | total |
| C | 4 | 4 | 4 | 12 |
| Bd | 0 | 4 | 0 | 4 |
| Rv | 0 | 0 | 4 | 4 |
| Ah | 4 | 0 | 0 | 4 |
| Bd & Ah | 0 | 4 | 4 | 8 |
| Ah & Rv | 4 | 4 | 0 | 8 |
| Rv & Bd | 4 | 0 | 4 | 8 |

**Table S4**. Number of individuals for each exposure group that showed signs of *Bactrachochytrium dendrobatidis* (Bd), Ranavirus (Rv), or both. Signs of Bd and Ranavirus were determined to be discoloration, sloughing, or thickening of the skin and hemorrhaging, respectively. Signs of disease in hosts were measured daily. Exposures are to Bd, Ranavirus, and *Aplectana hamatospicula* (Ah) either in sequential co-infections or in single infections. Number of individuals who showed signs of disease and experienced mortality are indicated by superscripted asterisks. No individuals who experienced mortality did not show signs of disease.

|  |  | **Signs** |  |
| --- | --- | --- | --- |
| **Exposures** | Bd | Ranavirus | Both |
| Bd & Ah | 10^*4^ | -- | -- |
| Ah & Bd | 1* | -- | -- |
| Rv & Bd | 0 | 4^*2^ | 0 |
| Bd & Rv | 2 | 4^*3^ | 5^*5^ |
| Rv & Ah | -- | 3^*2^ | -- |
| Ah & Rv | -- | 5^*1^ | -- |
| Rv | -- | 0 | -- |
| Bd | 0 | -- | -- |

**Fig. S1**. Growth rate (mm/week) of *Osteopilus* *septentrionalis* hosts: **A)** exposed to *Aplectana hamatospicula*, *Bactrachochytrium dendrobatidis* (Bd), or Ranavirus as single infections. Co-infections did not significantly change host growth rate (mm/week) when compared to single infections (see table S1 for Dunnett’s tests results). **B)** exposed to *A. hamatospicula*, Bd, or Ranavirus before co-infection with another parasite. Longer initial exposure time to Bd or Ranavirus before co-exposure to another parasite decreased host growth rate (Initial exposure time x initial parasite identity*:* F_2,136_=2.297, *p*<0.05; Time lag between infections (Bd): *F*_41,49_=5.00, *p*=0.03; Time lag between infections (Ranavirus): *F*_1,45_=6.617, *p*=0.013*)*. Data are shown as a conditional plot (*i.e*. controlling for everything else in the model), with the expected value (black hashed line), a 95% confidence interval or band for the expected value (gray band), and partial residuals (points) displayed.

**Fig. S2.** IgY antibody levels (OD values) of *Osteopilus* *septentrionalis* hosts that were not infected (C), were singly infected with *Aplectana hamatospicula* (Ah), *Bactrochochytrium dendrobatidis* (Bd), or Ranavirus (Rv), and were co-infected with *A. hamatospicula* and Ranavirus (AhRv), *A. hamatospicula* and Bd (AhBd), or Bd and Ranavirus (BdRv). Antibody levels were not significantly impacted by the identity and pairing of parasites that to which the hosts were exposed (Treatment group: *F*_6,35_=1.56, *p*=0.19) Data is shown as a conditional plot (*i.e*. controlling for everything else in the model), with the expected value (black hashed line), a 95% confidence interval for the expected value (gray band), and partial residuals (points) displayed.

**Fig. S3.** Log-transformed parasite load data over time for *O. septentrionalis* **A)** *Bactrachochytrium dendrobatidis (Bd)* loads on days 4, 8, 16, and 24 for host singly exposed Bd or initially exposed to Ranavirus (RvBd) or *Aplectana hamatospicula* (AhBd) before Bd exposure. Bd loads increased over time (χ²_3_ =162.15, *p*<0.001), but there was no effect of co-infection (χ²_2_ =1.23, *p*=0.54) or the interaction between time and co-infection on Bd load (χ²_6_ =9.08, *p*=0.17). **B)** Viral loads on days 2, 4, 8, and 24 for hosts singly infected with Ranavirus (Rv), or initially exposed to *A. hamatospicula* (AhRv) or Bd (BdRv) before Ranavirus exposure. Co-infected hosts had higher viral loads than singly infected hosts (χ²_2_ =12.17, *p*<0.01) and these differences changed through time (χ²_6_ = 20.66, *p*<0.001). Data are shown as a conditional plot (*i.e*. controlling for everything else in the model), with the expected value (black hashed line) and partial residuals (points) displayed.

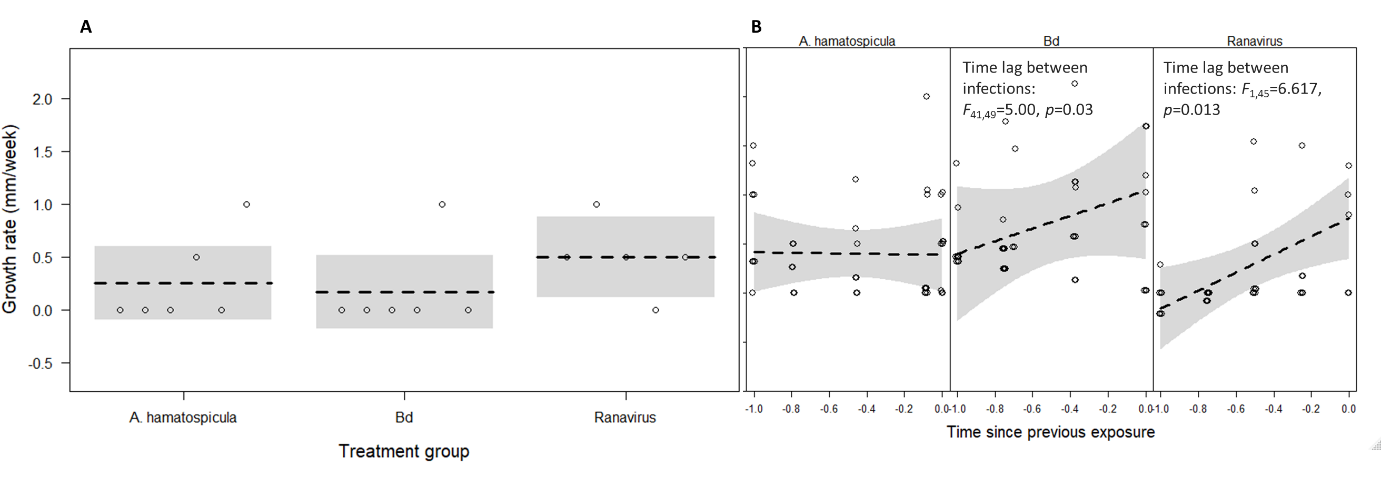

**Fig. S1**.

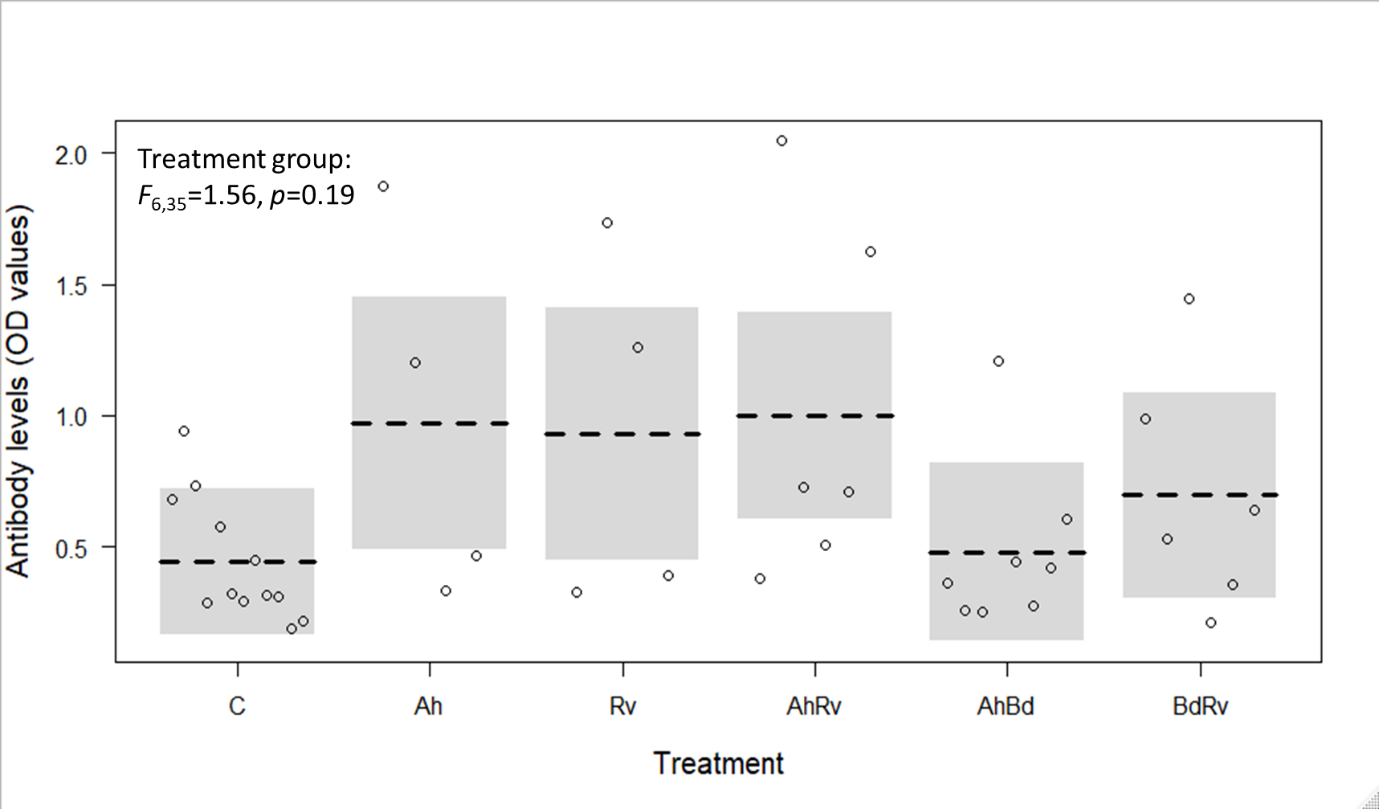

**Fig. S2.**

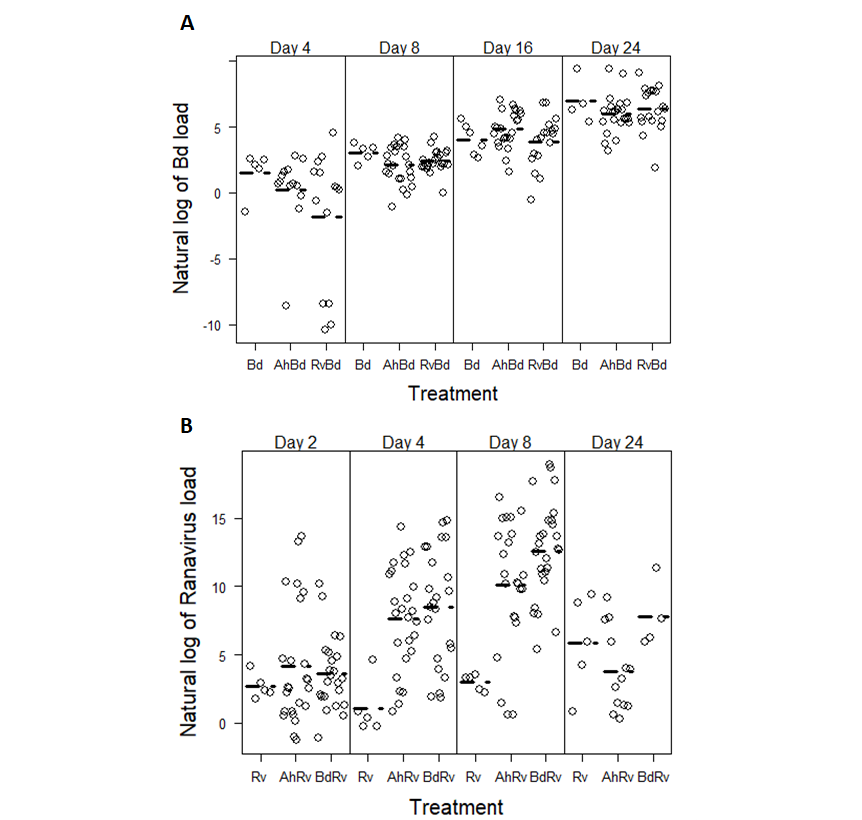

**Fig. S3**
